# Group size and activity influence shoal choice in Goldfish: Replication of zebrafish findings using a drift-diffusion model

**DOI:** 10.64898/2026.07.13.735552

**Authors:** Zhenbo Cheng, Jiangtao Ye, Hao Yan, Haohao Fu, Maoyu Wang, Xia Zhang, Mu Yuan

**Affiliations:** Zhejiang University of Technology, 288 Liuhe Road, Hangzhou, 310023, Zhejiang, China

**Keywords:** shoaling, group size, activity, drift–diffusion model, social decision-making, Goldfish

## Abstract

Animals often rely on social information when making movement decisions. In zebrafish, classic work showed that shoal size and shoal activity both bias shoal choice. Here we extend these effects in Goldfish (*Carassius auratus*) and extend them with a drift–diffusion model (DDM) account of individual evidence accumulation under dynamic social cues. Using a three-chamber linear arena, we quantified a focal fish’s position for 10 minutes while manipulating (i) numerical differences between flanking shoals and (ii) their activity (swimming speed) via temperature manipulation. ANOVA on time-proportion choices confirmed robust attraction to larger shoals; when shoal sizes were equal, the more active shoal was preferred. In combined manipulations, activity effects dominated at small numbers but saturated as group size increased, indicating a threshold-like integration where activity dominates at small shoal sizes (**<**3 fish) but saturates at larger sizes. We formalize these processes with a bounded DDM in which a sigmoidal stimulus function maps shoal size and average speed to momentary evidence, subject to random perturbations. The model reproduces the observed psychometric relations between relative numerosity, velocity differences, and choice. Our results (i) generalize zebrafish findings to a carp species with distinct ecology and physiology, and (ii) provide a compact mechanistic link between social cues and individual decision trajectories in dynamic social contexts.

## 1 Introduction

Grouping provides foraging and antipredator benefits across taxa. In fish, social information carried by *shoal size* and *shoal activity* (movement) strongly shapes individual choice [3–5]. A landmark study in zebrafish (*Danio rerio*) demonstrated that both the number of conspecifics and their activity bias shoal choice [1]. Numerous studies have since elaborated numerical discrimination in fishes [9–12] and collective decision rules [6–8]. Generally, fish prefer larger shoals (“numerical acuity”) to dilute predation risk, and prefer active shoals which may signal food availability or vigilance.

However, the decision-making *process*—how an individual continuously integrates dynamic social evidence over time—remains less understood. Most prior studies report only aggregate choice outcomes, ignoring the temporal dynamics. The Drift-Diffusion Model (DDM) offers a framework to bridge this gap by conceptualizing decisions as the accumulation of noisy evidence toward a threshold [2]. By applying DDM to shoaling behavior, we can test whether fish movement is merely a reaction to instantaneous cues or an integration of social evidence over time.

Furthermore, species differences matter for generalization. Zebrafish are small, diurnal cyprinids adapted to shallow, flowing habitats, with well-characterized visual circuits and standardized husbandry [13–16]. By contrast, Goldfish (*Carassius auratus*) are larger, more benthic-tolerant cyprinids with remarkable hypoxia tolerance and distinct ecological strategies [17, 18]. These differences motivate replication and modeling in *Carassius auratus*, testing whether previously reported shoal-size and activity effects extend to a carp with different sensory–physiological constraints.

Here we combine (i) an experimental extension in Goldfish that orthogonally manipulates shoal size and activity, analyzed with ANOVA as in behavioral work, and (ii) a DDM-based account in which a sigmoidal mapping from social cues (number *n* and average speed 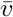) to stimulus 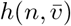 feeds a leaky, noisy integrator. We used the model to explain why fish do not always choose one side but instead move between both sides. Beyond corroborating zebrafish results [1], this synthesis supports a compact computational description of how fish translate dynamic social scenes into movement decisions.

## 2 Methods

### 2.1 Animals and Housing

Goldfish (*Carassius auratus*) were sourced from a local aquaculture supplier in Hangzhou, China. They were housed in 300 L stock tanks at 25^°^C using dechlorinated tap water, under a 12:12 h light:dark cycle, and fed commercial flake food twice daily. A total of approximately 200 fish were used in the study. All subjects were naive to the experimental apparatus prior to testing.

### 2.2 Apparatus

The experimental setup consisted of a three-chamber linear tank (46 cm × 18 cm × 20 cm) with a water depth of 12 cm (Fig. 1). Two transparent partitions separated the tank into three compartments: a central observation chamber (16 cm length) for the focal fish, and two end stimulus chambers (15 cm length each) for the stimulus shoals. The outer walls were opaque to prevent external visual distractions. The central chamber was virtually divided into five zones (*k* ∈ {−2, −1, 0, 1, 2}) for analysis (Fig. 2). Behavior was recorded using a top-down CMOS camera at 30 fps.

**Fig. 1.**
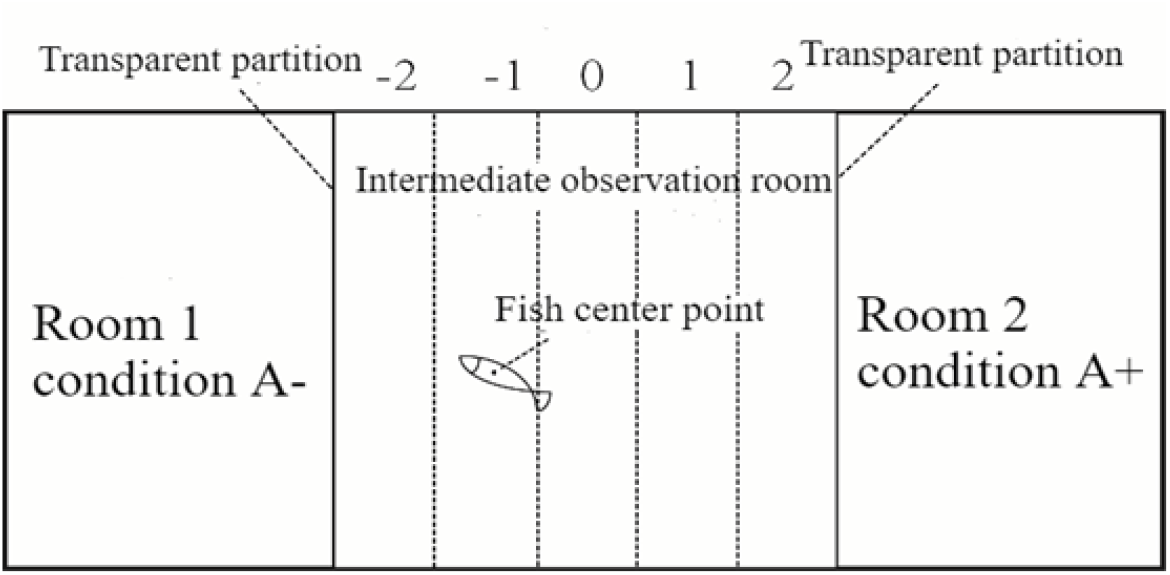
Top view of the three-chamber linear arena (opaque outer walls, transparent end chambers).

**Fig. 2.**
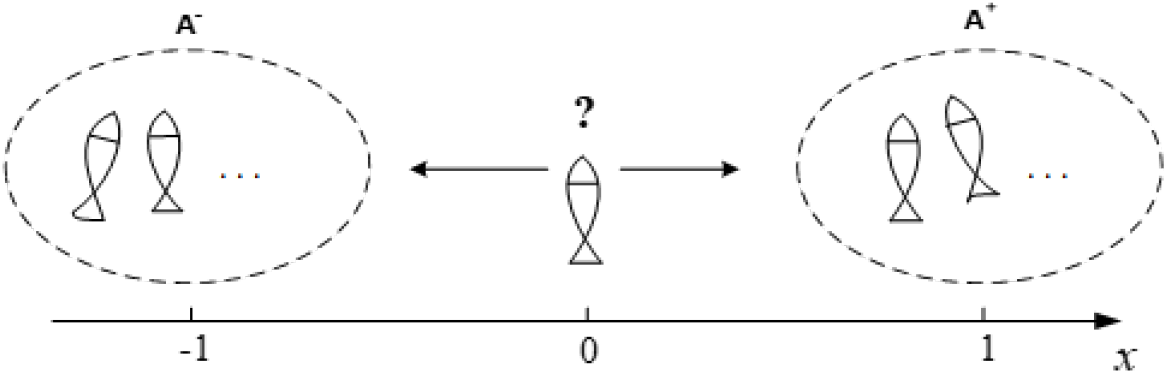
One-dimensional abstraction of the choice geometry with options *A*^±^ at ±1 and zones *k* ∈ {−2, −1, 0, 1, 2}.

### 2.3 Experimental Conditions and Activity Manipulation

We conducted three types of choice experiments: (i) Size Asymmetry: Varying the number of fish (1 to 6) on one side while keeping the other constant; (ii) Activity Asymmetry: Keeping group sizes equal but manipulating swimming speed; and (iii) Combined Manipulation: Varying both size and activity simultaneously (e.g., a small active shoal vs. a large inactive shoal).

To manipulate activity levels (swimming speed) without using invasive drugs, we controlled the water temperature in the end stimulus chambers only. The baseline temperature was 25^°^*C*. To increase activity, the temperature in a specific end chamber was elevated to 30^°^*C*; to decrease activity, it was lowered to 20^°^*C*. Temperature was maintained using external submersible heaters/coolers that did not obstruct the visual field. Crucially, the water in the central observation chamber remained strictly at 25^°^*C* throughout all trials to ensure the focal fish’s metabolism and locomotion were not thermally affected. Activity levels were validated in pilot trials: fish at 30^°^*C* exhibited significantly higher average speeds than those at 25^°^*C*, while fish at 20^°^*C* were slower.

### 2.4 Procedure and Randomization

To eliminate any potential side bias (preference for the physical left or right side of the room), the presentation of the stimulus was fully randomized. For any given condition (e.g., “Large Shoal” vs. “Small Shoal”), the “target” stimulus (e.g., the larger shoal) was presented on the left side for half of the trials (*n* = 10) and on the right side for the other half (*n* = 10), in a randomized sequence. In each trial, stimulus fish were allowed to acclimatize in the end chambers for 5 minutes. A focal fish was then introduced to the center of the middle chamber. Recording began after a 5-minute acclimation period and lasted for 10 minutes. Each fish was used only once as a focal subject.

### 2.5 Data Analysis and Statistics

Video data were analyzed to extract the centroid of the focal fish frame-by-frame. The primary metric was the Proportion of Time (*P*) spent in the preference zones (*k* = ± 2) adjacent to the stimulus chambers. Data Transformation for Analysis: To facilitate clear reporting, data from randomized trials were aligned logically rather than physically. We designated “Side A-” as the side with the higher magnitude of the manipulated variable (i.e., larger number or higher activity), regardless of whether it was physically on the left or right. Conversely, “Side A+” refers to the alternative side.

We used one-sample t-tests to compare the proportions *r*_±2_ (time spent near each end chamber) and side-preference ratios under each manipulation to the expected mean of 0.5 (arcsine transformed), which would occur if the fish randomly allocated their time across the two chambers. A significant difference indicated a preference for one of the chambers under the respective manipulation. The overall effects of group size and activity on preference score and total time spent in each chamber were analyzed using ANOVA. All data are presented as mean ± standard error of the mean (SEM).

## 3 Results

### 3.1 Preference for larger shoals

In experiments manipulating shoal size asymmetry, we compared shoals ranging from equal sizes (1:1) to highly asymmetric (6:1), with the larger shoal on one side (side assignment randomized and balanced across trials to control for bias, as described in Methods). Focal Goldfish rarely dwelled centrally and instead oscillated between ends of the central lane. Time-proportion near each side (regions *k* = ±2; Fig. 1 & 2) increased with the number of adjacent conspecifics, supporting the concept of mutual attraction among conspecifics (Fig. 3). The fish displayed a clear preference for being near the side walls, particularly in regions *k* = −2 or *k* = 2, when the numbers of individuals on either side were unequal. Minimal time was spent in the central, equidistant region (*k* = 0).

**Fig. 3.**
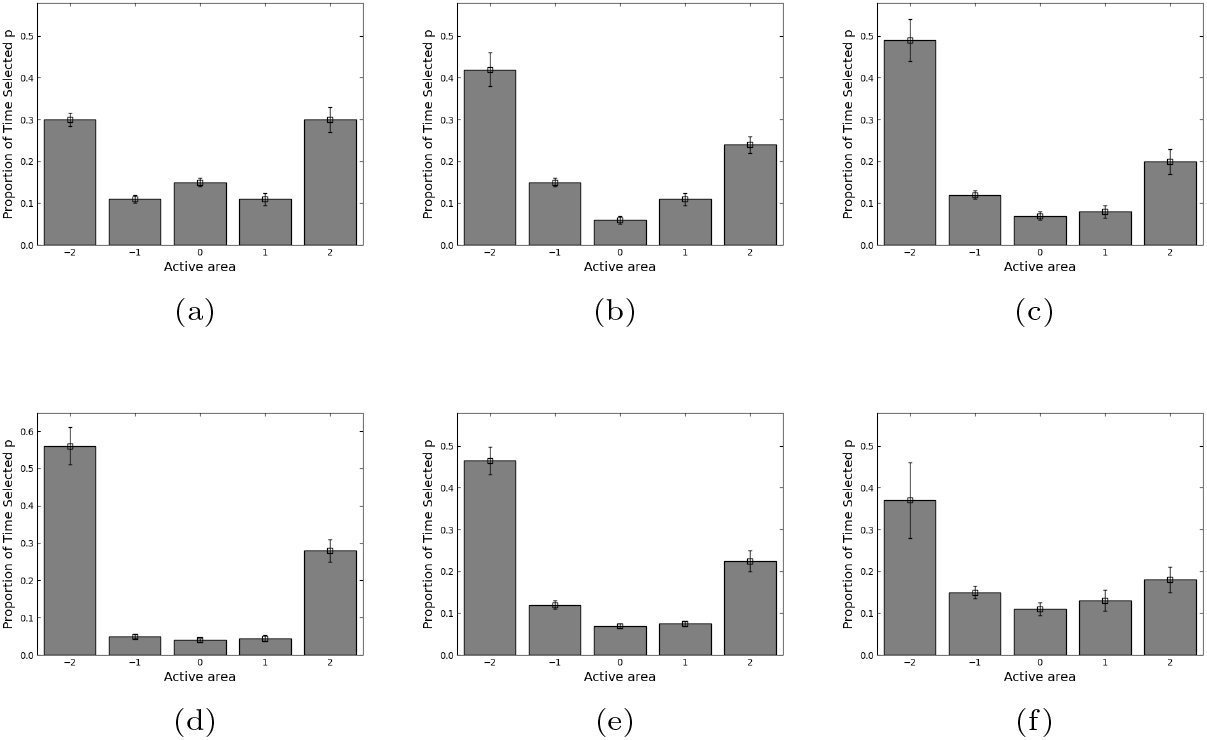
Time-distribution across zones as numerosity on one side increases (panels correspond to 1:1 through 6:1 comparisons).

When both sides had one fish (1:1), end preferences were statistically indistinguishable (*F* (1, 18) = 5.51 × 10^−4^, *p* = 0.9816 > 0.05). As the left-side number increased (from 1:1 to 6:1), preference for that side rose and saturated for 4–6 fish (Fig. 4). ANOVA showed a significant step between 2 and 3 (*F* (1, 18) = 7.93, *p* = 0.0124 < 0.05), but no significant differences among 3–6 (*F* (3, 36) = 1.2, *p* = 0.3368 > 0.05), indicating a perceptual/decision threshold beyond which additional individuals do not enhance stimulus.

**Fig. 4.**
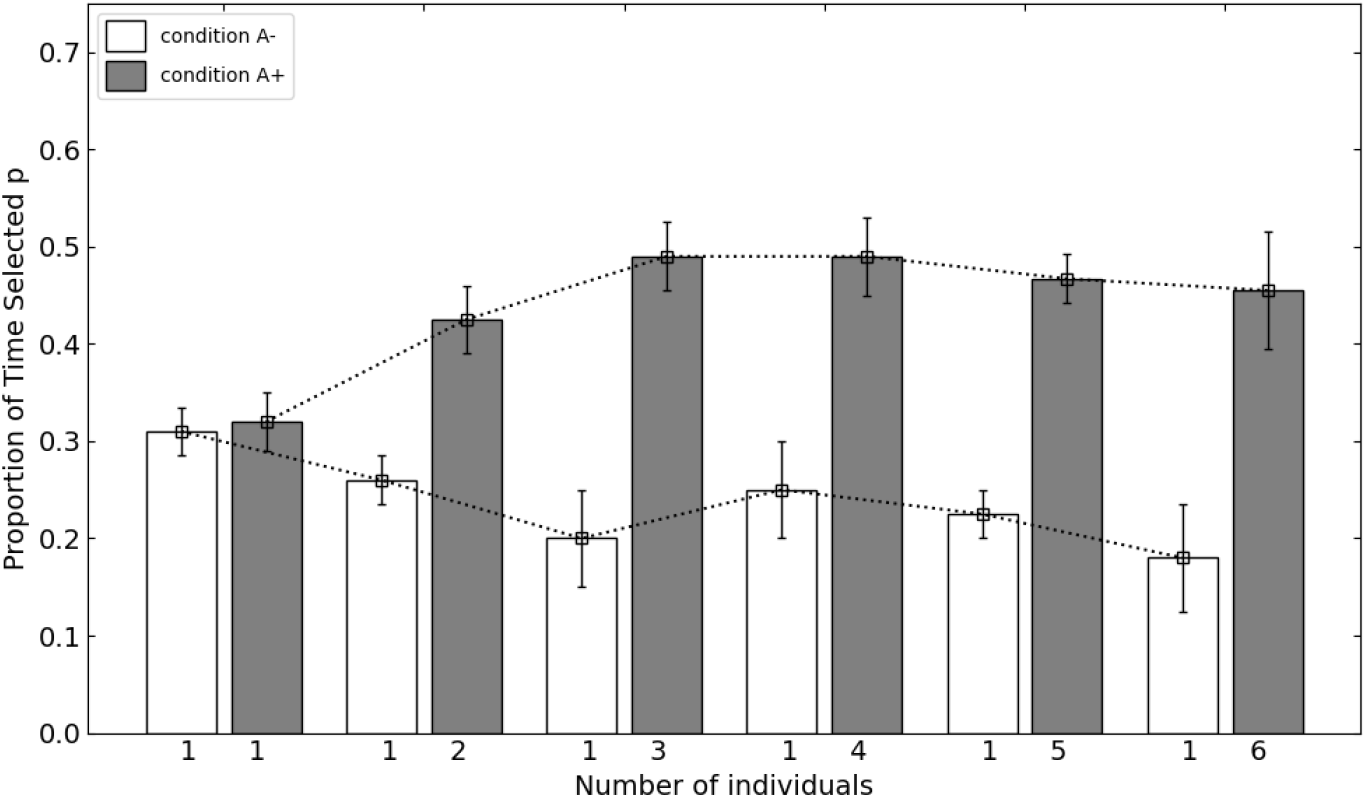
Proportion of time spent choosing a side vs. group-size difference. Saturation appears around 4–6 individuals.

The difference in group size Δ*N* was defined as 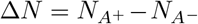, where 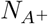 and 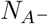 are the numbers of individuals on the respective sides. Disparities in individual counts led to significantly different probabilities, with the proportion of time spent near the larger shoal increasing as Δ*N* grew, stabilizing around four to six individuals.

### 3.2 Activity biases choice when sizes are equal

In experiments where shoal sizes were equal on both sides (ranging from 1 vs. 1 to 6 vs. 6 fish), we manipulated activity by altering temperature in one end chamber (±5^°^C from ambient 25^°^C), either elevating temperature to increase activity or reducing it to decrease activity on one side (randomized assignment). Two protocols were used: (1) reducing activity on one side via cooling, while the other remained at ambient temperature; (2) elevating activity on one side via heating, while the other remained normal. Each condition was tested in 10 groups per protocol, totaling 20 trials per group size.

When both sides contained equal numbers, the more active shoal was preferred (Fig. 5). Across 6 groups (120 trials), the proportion choosing a side increased with that side’s average swimming speed. The normal swimming speed ranged between 1.15 cm/s and 2 cm/s; speeds below this reduced selection probability, while higher speeds increased it. There was a significant difference in selection behavior under different activity levels (*F* (1, 39) = 23.67, *p* = 1.82 × 10^−5^ < 0.01).

**Fig. 5.**
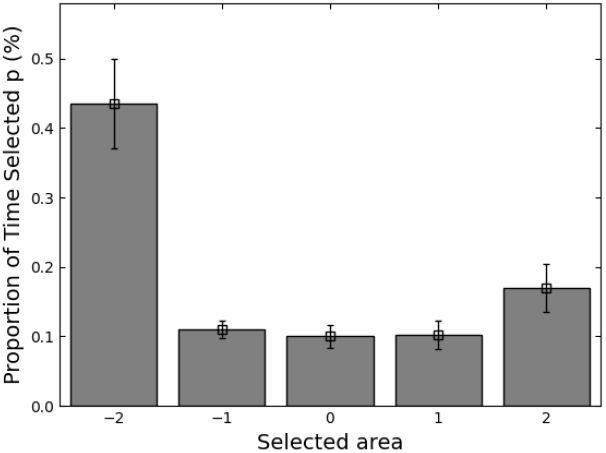
Time distribution under activity manipulations with equal group sizes; more active side is preferred.

The average velocity under each condition was calculated, and a regression equation was fitted: *p* = 0.1316*v* + 0.122, where *p* is the selection proportion and *v* is average velocity, indicating a positive relationship between activity and preference.

With 1 vs. 1 and 2 vs. 2, preferences differed significantly in favor of the faster side (*F* (1, 38) = 23.67, *p* < 0.05; *F* (1, 38) = 4.78, *p* < 0.05). As numerosity increased (≥ 3 per side), activity-based differences diminished and were no longer significant (*F* (1, 38) = 0.12, *p* = 0.7334 > 0.05) (Fig. 6 & 7). The fish spent a higher proportion of time near the more active group, supporting the hypothesis that activity levels influence preference when sizes are equal.

**Fig. 6.**
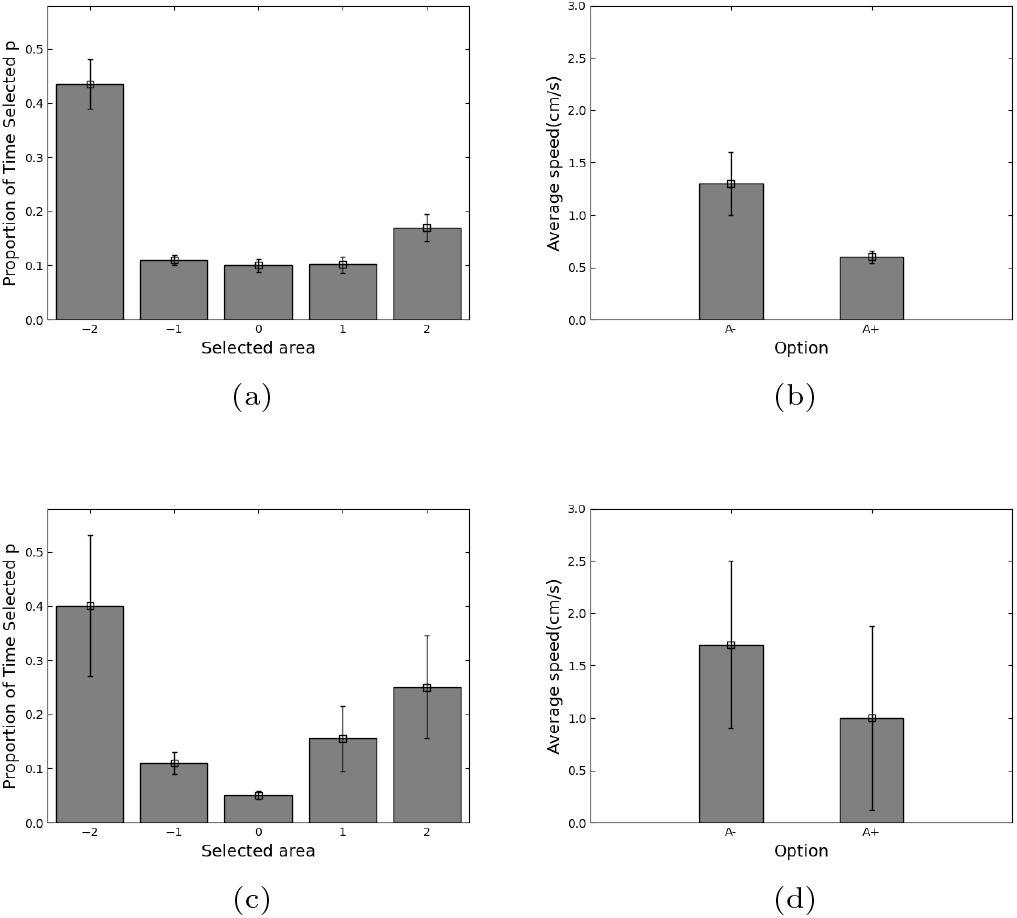
Equal-size conditions (1 vs. 1; 2 vs. 2): choice proportions and speed distributions across the two activity protocols (temperature elevation vs. reduction).

**Fig. 7.**
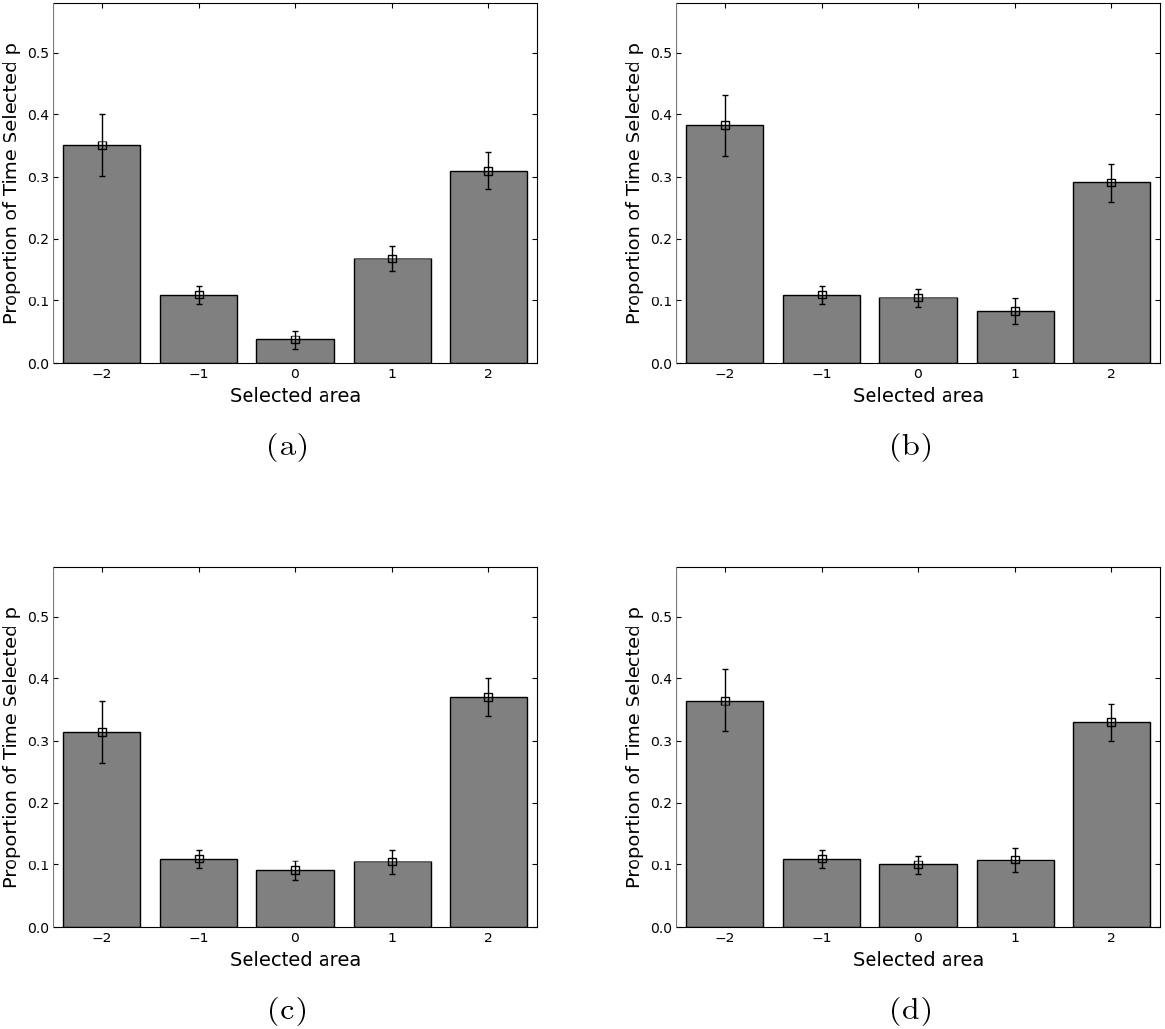
Equal-size conditions (3–6 per side): activity effects diminish as numerosity increases.

### 3.3 Number–activity integration: dominance at small n and saturation at larger n

In combined manipulations, we orthogonally varied shoal size and activity, for example, by elevating activity (via +5^°^C) on the numerically smaller shoal (e.g., 1 vs. 2, 2 vs. 3) or reducing it on the larger one, across small (1-3 fish) and large (4-6 fish) absolute sizes. This replicated the size asymmetry experiments but with the modification of increasing activity on the smaller side via heating to examine if higher activity could alter preference.

Combining manipulations showed that elevating activity on the numerically smaller side increased selection of that side when absolute numbers were small (Fig. 8a), but this advantage waned as group size grew (Fig. 8b). The difference in selection proportion diminished with increasing group size, indicating that in smaller groups, activity has a significant impact, but as size increases, group convergence reduces its influence.

**Fig. 8.**
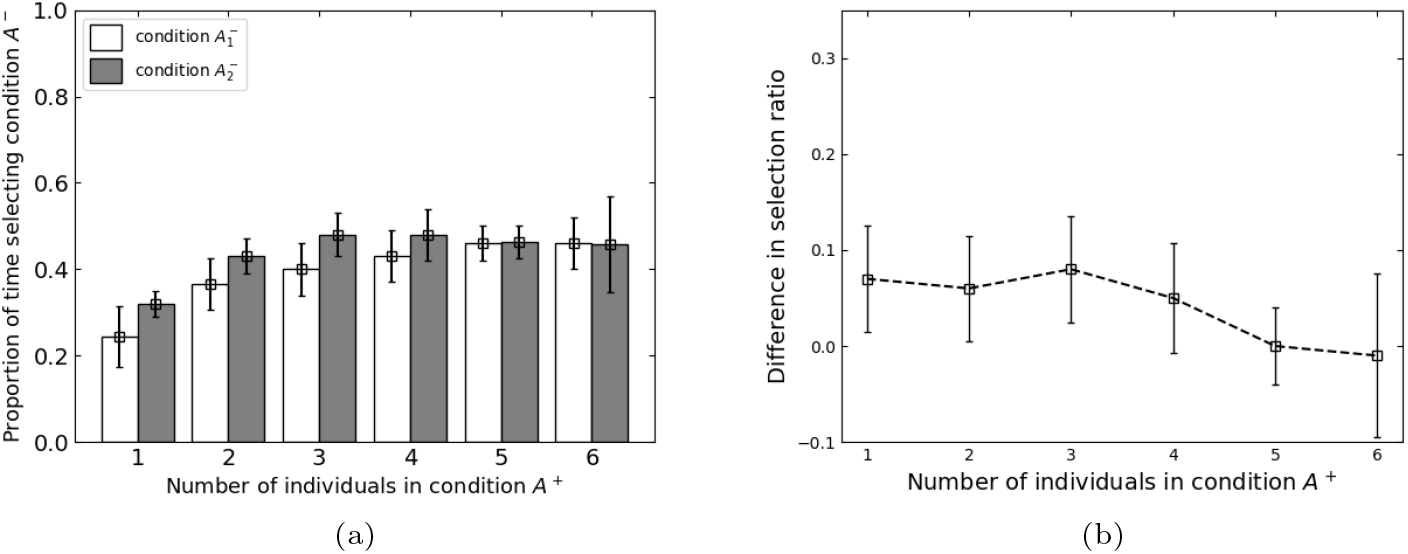
Combined manipulations: (a) elevating activity on the numerically smaller side increases selection at small *n*; (b) effect wanes as *n* grows.

The experimental results support that when activity of the smaller group is increased, there is a corresponding rise in the proportion selecting that side. However, with larger groups, decisions align more with overall group behavior, leading to reduced differences in selection proportions.

## 4 Modeling individual decisions with a DDM

The Drift-Diffusion Model (DDM) [2] is a well-established framework for modeling decision-making under uncertainty, where individuals accumulate evidence over time to reach a threshold and make a choice. In the context of shoal choice in fish, this model is particularly useful for understanding how an individual integrates dynamic social cues, such as shoal size and activity, when making a decision.

In our study, the DDM was applied to formalize how Goldfish accumulate evidence based on the social information provided by surrounding shoals. Specifically, the model posits that decision-making is driven by a process of bounded evidence accumulation where the fish integrate both numerosity (shoal size) and activity (average swimming speed) as distinct cues. The evidence is accumulated over time, influenced by random perturbations and social cues, until a threshold is reached, leading to a choice of one shoal over the other.

We chose to model the decision-making process using a DDM for several reasons: (i) Evidence Accumulation: The decision to select a shoal likely involves accumulating evidence over time. The fish are exposed to fluctuating cues (e.g., numerosity and speed) that influence their choice. The DDM provides a natural framework for this process, where each stimulus (shoal size, activity) contributes to a dynamic accumulation of evidence that is ultimately compared to a threshold.

(ii) Noise and Stochasticity: Real-world decision-making often involves noise or randomness, as observed in the oscillatory shuttling behavior of the fish in the arena. The DDM incorporates this noise by modeling random fluctuations in the decision process. This is particularly useful in capturing the observed behavioral patterns, where the fish switch between shoals due to the random perturbations inherent in the environment.

(iii) Psychometric Functions and Thresholding: Our empirical results suggest that the influence of group size saturates beyond a certain threshold, and the effects of activity diminish as group size increases. The DDM allows us to model this threshold-like behavior, where the integration of activity effects dominates at smaller group sizes but saturates at larger ones. This feature of the DDM aligns well with the observed data.

(iv) Modeling Decision Trajectories: The DDM not only allows us to capture the final choice but also provides insight into the underlying decision trajectory. By simulating the continuous process of evidence accumulation, we can investigate how the fish arrive at their decisions over time, which helps us understand the temporal dynamics of social decision-making.

We implemented stimulus integration in the DDM using a sigmoidal function 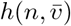 that maps shoal size (*n*) and average speed 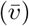 to a momentary evidence signal. The function is defined as:

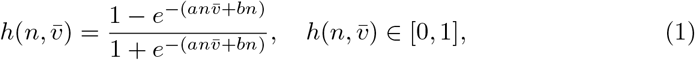

where *a* and *b* are parameters that control the sensitivity to activity and numerosity, respectively. This sigmoidal function captures the idea that the influence of both cues saturates at higher values, reflecting the diminishing returns observed in our empirical data.

When *a* = 0.8, and *b* = 0.2 are set in equation (1), the resulting sigmoidal mapping 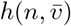 is illustrated in Fig. 9. This function effectively captures the combined influence of shoal size and activity on the stimulus strength, which is then used as input to the DDM for evidence accumulation.

**Fig. 9.**
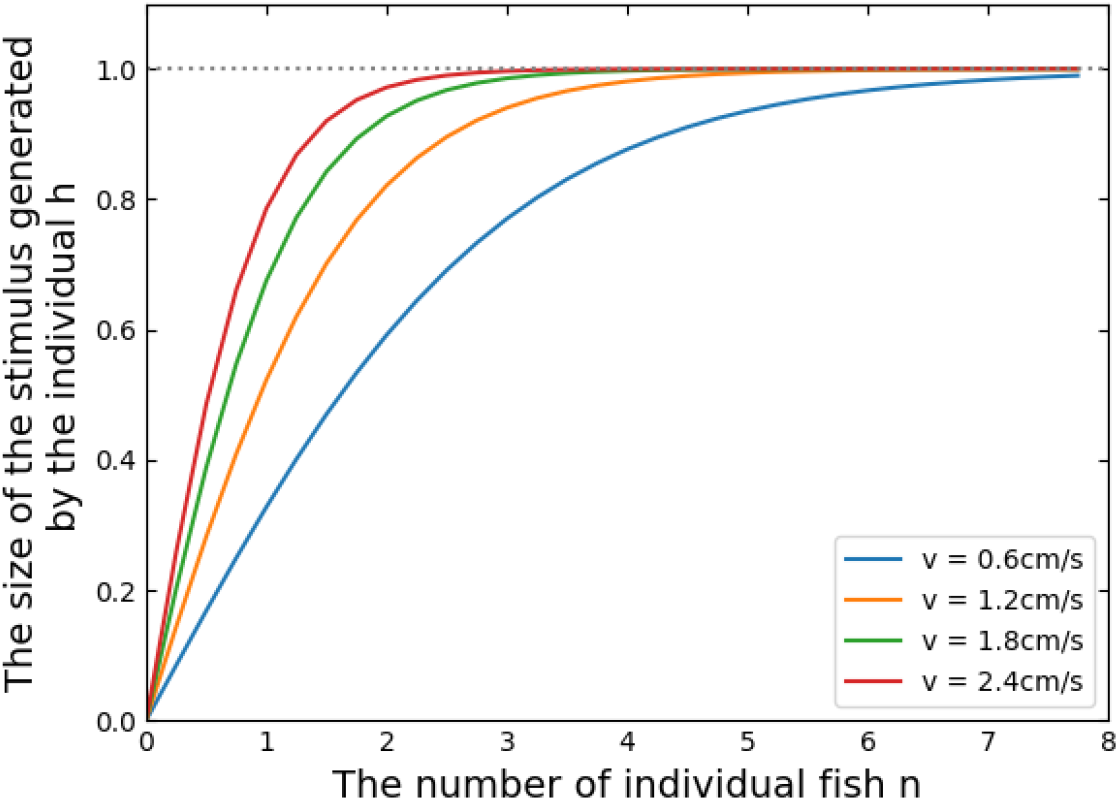
Sigmoidal mapping 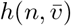 from numerosity (*n*) and activity 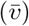 to stimulus strength (example parameters *a*=0.8, *b*=0.2).

To incorporate the effect of random environmental perturbations on stimulus perception, we introduce the random variable *H*_*t*_, which represents the stimulus generated at each discrete time step. The variable *H*_*t*_ is defined as a Bernoulli random variable:

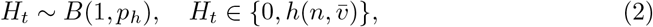

where *p*_*h*_ is the probability of perceiving the stimulus 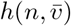 at time *t*. This formulation captures the idea that the fish may not always perceive the social cues perfectly due to environmental noise or other factors.

Let *x*_*t*_ be the position of the fish at time *t*. The position *x*_*t*_ is normalized to the interval [−1, 1], where −1 corresponds to the leftmost position (near the left shoal), 1 corresponds to the rightmost position (near the right shoal), and 0 represents the central position in the arena. The fish’s movement is influenced by the perceived stimuli from both shoals, denoted as 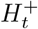 for the right shoal and 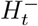 for the left shoal.

The evidence accumulation process in the DDM is then modeled as:

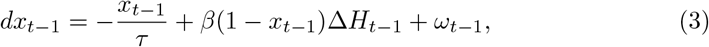

where 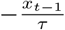 represents the temporal decay of the influence of the previous position, *β*(1 − *x*_*t*−1_)Δ*H*_*t*−1_ captures the influence of the current stimulus difference on the decision variable, and *ω*_*t*−1_ is a Gaussian noise term with mean 0 and variance *σ*^2^. The parameter *τ* controls the rate of decay, while *β* scales the influence of the stimulus.

Δ*H*_*t*−1_ defined as:

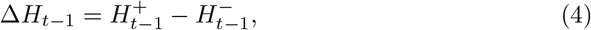

represents the difference in perceived stimuli from the two shoals at time *t* −1.

The position of the fish at time *t* is then updated as:

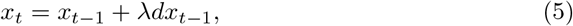

where *λ* is a scaling factor that translates the accumulated evidence into a positional change.

Considering that *x*_*t*_ ∈ [−1,1], if *x*_*t*_ > 1 or *x*_*t*_ < −1 occurs, it indicates that the fish has exceeded its parallel distance from adjacent individuals and entered their personal space. To avoid collisions from group pressure, the individual will stay within the boundary (i.e., *x*_*t*_ = ±1). Thus, the position of the individual fish at the next moment *x*_*t*_ is given by:

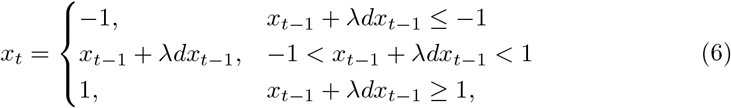

To capture the stochastic nature of Goldfish decision-making, we incorporate Gaussian noise *ω*_*t*−1_ ~ *N*(0, *σ*^2^) into the evidence accumulation process (Eq. 3). This noise term serves several purposes. First, it models random environmental and internal perturbations, such as water flow variations, visual occlusions, or attentional fluctuations, which may disrupt the focal fish’s perception of shoal size (*n*) and activity 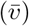. Second, Gaussian noise accounts for the observed oscillatory shuttling between shoals (Fig. 10a), preventing overly deterministic predictions and reproducing empirical trajectories (Fig. 10b). Finally, the noise complements the Bernoulli-distributed stimulus perception (Eq. 2), together capturing both discrete detection and continuous integration of social cues. The variance *σ*^2^ parameterizes the noise strength, enabling the model to fit varying degrees of behavioral stochasticity across experimental conditions.

**Fig. 10.**
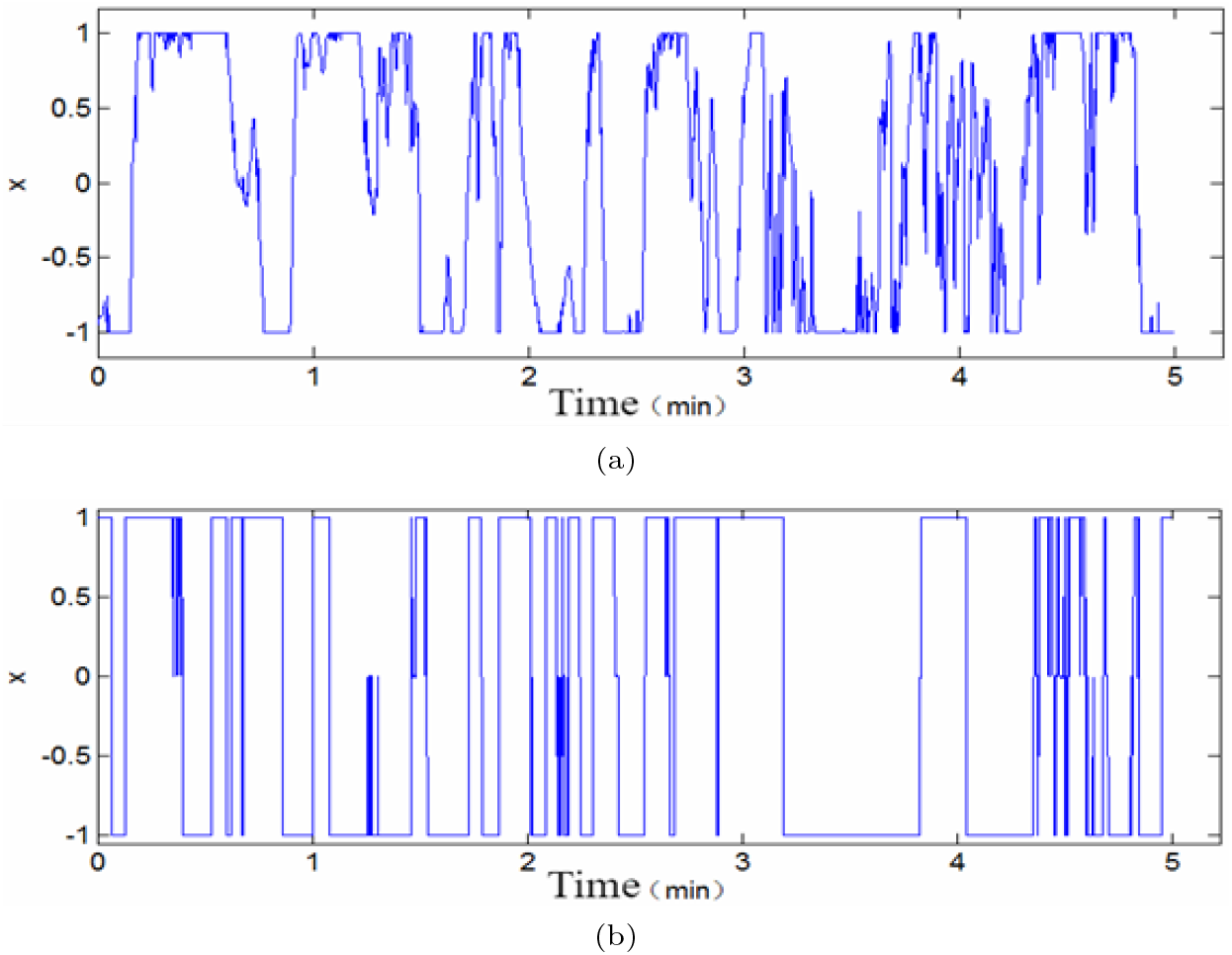
(a) Empirical position *x*_*t*_ over time for an equal-size, equal-activity condition. (b) Model simulation reproducing oscillatory shuttling.

This is a fish individual decision-making model based on the DDM framework, which captures the dynamics of evidence accumulation influenced by social cues and random perturbations. The model parameters can be adjusted to fit empirical data, allowing for a quantitative understanding of how fish integrate shoal size and activity when making movement decisions. Fig. 10a shows an empirical trajectory of a focal fish in an equal-size, equal-activity condition, exhibiting oscillatory shuttling between the two shoals. Fig. 10b presents a simulation of the DDM model that reproduces this oscillatory behavior, demonstrating the model’s ability to capture the dynamics of fish movement decisions in response to social cues.

## 5 Discussion

Our study successfully replicates and extends the classic findings of shoal choice in zebrafish to Goldfish (*Carassius auratus*). We demonstrate that both numerical quantity and activity level are critical determinants of social decision-making, yet they operate through a nonlinear integration rule. Specifically, we found that preference for larger shoals follows a psychometric function that saturates at approximately 3–4 individuals [9, 11], and that high activity acts as a potent attractant that can override numerical inferiority, particularly in small groups [1, 3].

### 5.1 The Trade-off Between Quantity and Quality

The observation that activity exerts a dominant influence at small group sizes but saturates as numerosity increases suggests a specific ecological trade-off. In small shoals, the “dilution effect” is weak, making individual predation risk high. In this context, an active neighbor may serve as a high-value signal of vigilance or foraging success (“quality”), offering immediate benefits that outweigh the lack of numbers [5]. However, as group size increases, the safety benefits from the dilution effect arguably become the primary driver, rendering the marginal benefit of additional activity less significant. This interpretation challenges the view that activity is merely a visual proxy for group size (i.e., making a group appear larger); rather, our data suggests it is a distinct social cue whose weight is dynamically adjusted based on the current context [7]. The slightly higher saturation threshold observed in goldfish (3–4) compared to zebrafish (2–3) [1] may reflect species-specific differences in visual acuity or social aggregation tendencies adapted to their respective benthic versus pelagic habitats [15, 17].

### 5.2 Mechanistic Insights from the Drift-Diffusion Model

Beyond descriptive behavioral patterns, our application of the Drift-Diffusion Model (DDM) provides a mechanistic explanation for *how* these decisions unfold over time. Previous studies often treated shoal choice as a static outcome [10]. In contrast, our DDM formalizes the decision as a continuous, noisy evidence accumulation process [2]. Crucially, the model offers two key insights: (1) **Sensory Saturation:** The sigmoidal stimulus function 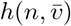 quantitatively captures the psychophysical limits of the fish’s sensory system, confirming that the behavioral plateau is a perceptual constraint (Weber’s Law) rather than a motor limitation. (2) **Oscillatory Dynamics:** The model explains the frequent “shuttling” behavior observed in our experiments not as random indecision, but as a signature of a leaky integrator interacting with stochastic noise. This suggests that “choosing” a shoal is not a one-off event but a dynamic maintenance of social proximity driven by fluctuating evidence [4].

### 5.3 Limitations

We acknowledge certain limitations. First, while we strictly controlled the temperature of the central observation zone to isolate visual cues, temperature-induced metabolic changes in the stimulus fish could theoretically introduce non-visual chemical cues, although our flow-through setup minimized this risk [13]. Second, the linear arena constrains natural 3D shoaling dynamics [8]. Future work employing high-resolution 3D tracking and neural recording could further validate the DDM’s accumulation variable against specific brain circuit activities [16].

## 6 Conclusion

In summary, this study generalizes the rules of social decision-making from zebrafish to goldfish, highlighting a conserved algorithm where social cues are weighted nonlinearly: activity dominates when numbers are scarce, while numerosity prevails as groups grow. By formalizing this process with a Drift-Diffusion Model, we bridge the gap between ethological observation and computational mechanism. This framework not only explains the static preferences reported in prior literature but also accounts for the dynamic trajectories of individual animals, offering a robust tool for comparative behavioral neuroscience.

## Declarations

### Compliance with Ethical Standards

The authors confirm that the manuscript adheres to the ethical standards of the journal, including originality, avoidance of plagiarism, and proper citation. All procedures were conducted in accordance with relevant guidelines for animal research.

### Ethical Approval

All applicable international, national, and institutional guidelines for the care and use of animals were followed. This study complied with local regulations for non-invasive behavioral observations on fish, and formal ethical approval was not required as per institutional policies at Zhejiang University of Technology. A total of approximately 200 goldfish were used, and efforts were made to minimize suffering through proper husbandry and temperature manipulations.

### Informed Consent

Not applicable, as this study did not involve human participants.

### Author Contributions

Z.C. and J.Y. conceived the idea and designed the experiments. Z.C., J.Y. and H.F. performed the experiments and analyzed the data.

M.W. and H.Y. contributed to data visualization and prepared the figures. M.Y. and X.Z. provided supervision and guided the research. Z.C. and J.Y. wrote the main manuscript text. All authors reviewed the manuscript.

### Data and Code Availability

The datasets and code supporting the findings of this study are available from the corresponding author upon reasonable request. The simulation scripts include Python and MATLAB implementations of the drift-diffusion model used to simulate fish decision-making.

